# Detection of a novel African-lineage-like Zika virus naturally infecting free-living neotropical primates in Southern Brazil

**DOI:** 10.1101/828871

**Authors:** Paula Rodrigues de Almeida, Luiza Presser Ehlers, Meriane Demoliner, Ana Karolina Antunes Eisen, Viviane Girardi, Cíntia De Lorenzo, Matheus Viezzer Bianchi, Lauren Mello, Saulo Pettinati Pavarini, David Driemeier, Luciana Sonne, Fernando Rosado Spilki

## Abstract

Mosquito borne flaviviruses cause a series of important diseases in humans and animals. These viruses are maintained in cycles involving replication in mosquito and in vertebrate hosts. Most natural hosts are vertebrate animals living in sylvatic or peridomestic environments. Human contact with these environments may result in host shifts that lead to the establishment of urban transmission cycles. Zika virus is a *Flavivirus* that persists in nature in a transmission cycle involving non-human primates (NHP). Its recent emergence in Brazil has shed light upon the importance of surveying this agent in Brazilian sylvatic environments. Here we present histopathological and molecular evidence that free ranging howler monkeys (*Alouatta guariba*) in Southern Brazil are infected by ZIKV closely related to African lineage MR766. Nine NHP were nested RT-PCR positive for ZIKV RNA. Sequence analysis revealed 96 to 98% identity to ZIKV MR766 and 85% identity to ZIKV P6-740, the current epidemic strain. The affected howler monkeys presented discrete inflammatory infiltrates in several tissues and immunohistochemichal (IHC) labeling of viral antigen was observed in placenta. These findings point to the circulation of African lineage Zika virus in the Americas in non-human primates. And raises the possibility that ZIKV was introduced into the Americas on more than one occasion.

## Introduction

Arboviral diseases represent a burden of up to 20% all infectious diseases in the World, and many arbovirus diseases are zoonotic^1,2^. The majority of zoonotic emerging infectious diseases are acquired from mammals, and non-human primates (NHP) are among the most common primary sources of novel viruses for human beings. External pressures such as urbanization and agricultural expansion impact the host populations, can causing spillover events^2^. Zika virus (*Flaviviridae*, genus *Flavivirus*) was discovered in the Zika Forest in Uganda, in 1947. The lineage of ZIKV discovered emerged in Uganda between 1892 and 1943, probably around 1920. Currently, ZIKV is distinguished in three different lineages, the East African, West African and the Asian/American ^3^.

ZIKV is highly adapted to sylvatic environments transmitted among NHP by forest dwelling mosquitoes^4^. The detection of ZIKV in Asia related to major outbreaks and Guillain Barré Syndrome, and subsequently in the Americas causing congenital microcephaly highlighted the possibility of a sylvatic cycle establishment ^5^. African lineages of ZIKV have never been detected in the Americas, these lineages of ZIKV were only isolated from mosquito pools and NHP samples, while naturally occurring ZIKV from Asian lineage was mostly isolated from human samples and mosquitoes^6^. However, the circulation of epidemic strain of ZIKV in capuchin monkeys (*Sapajus apella*) and marmosets (*Callithrix penicillata*) was evidenced in Brazil^2,7,8^.

The recent ZIKV emergence in Yap Island causing Guillain Barré Syndrome and in Brazil causing thousands of microcephaly cases highlighted the variety of possible clinical and pathological outcomes caused by this virus ^9,10,11^. ZIKV infection clinical outcome varies, and it can manifest as a Dengue-like disease, with mild fever and rash, a life-threatening Guillain-Barré syndrome, and also several congenital neurological malformations, including microcephaly that are now recognized as congenital zika syndrome^9,10,11,12^.

Unlike most viral infections, Asian ZIKV lineages are capable of pass through placental barrier and reach the fetus in pregnant NHP^13^. Pathological findings in human fetuses infected with ZIKV include thickening of leptomeninges, subcortical calcifications with cortical displacement, ventricular derangement, astrocytosis, macrophagic infiltrate and microglial proliferation^10^. ZIKV infection in mice causes apoptosis, lymphocytic infiltrate and persistent infection in glomerular and tubular cells, moreover, ZIKV lineage MR 766 produces higher viremia and cytopathic effects in kidney cells than the Asian lineage ^14^. Inoculation of Asian ZIKV in Rhesus monkey spread to several tissues including kidney, bladder, uterus, with persistence in central nervous tissue (CNS), joints and lymphnodes. Histologically, in ZIKV experimentally infected animals, inflammatory infiltrates were observed in multiple tissues ^15,16^. Viral persistence in experimentally infected NHP has been already detected in liver, spleen, kidney, uterus, CNS, bladder, testes, heart, lymph nodes, joints, and bladder ^15,16^.

Here we report clinical and pathological findings of free ranging howler monkeys (*Alouatta guariba*) naturally infected with a ZIKV. The strain detected from nine animals was phylogenetically closer to MR-766 lineage of ZIKV than from the currently epidemic strains circulating in Brazil, the tenth case, a neonate howler monkey, was positive for ZIKV and presented several pathological lesions compatible with Congenital Zika Syndrome (CZS). Furthermore, molecular characterization and phylogenetic classification of the virus detected in the first nine cases are described. Our findings indicate that an african-related lineage of ZIKV is circulating in wildlife in the Americas and imply that the ZIKV lineage reported here may be co-circulating with the epidemic strain under the limits of detection and characterization, possibly as a consequence of the high levels of circulation of the Asian/American lineage in the Americas; also, the absence of detection of this strain could be attributable to the region and habitat where it was detected – sylvatic areas of a temperate to subtropical state of Brazil, with low prevalence of arboviral diseases.

## Results

### nested RT-PCR positive NHP

A total of 128 tissue samples belonging to 50 NHP were screened through nested RT-PCR for Flavivirus. Nine NHP were positive for Flavivirus in the nested RT-PCR test (Table 1). All positive NHP involved in the outbreak were *Allouata guariba* (howler monkey, Figure 1). Four howler were from Porto Alegre city (30.0346°S, 51.2177°W),1 was found in Camaquã city (30.8494°S, 51.8048°W), 1 was found in Chuvisca city (30.7564°S, 51.9787°W) and 1 was found in Triunfo city (29.9381°S, 51.7153°W), two NHP were from the metropolitan region of Porto Alegre, however we did not have their location informed. Positive NHP locations were connected either by fragments of Atlantic forest or by water of the Guaíba and Jacuí rivers (Figure 2).

**Table 1.** Nested PCR positive NHP

|  | Tissue |  |  |  |  |
| --- | --- | --- | --- | --- | --- |
|  | Brain | Cerebellum | Spinal Cord | Liver | Placenta |
| NHP1 | (-) | NT | NT | NT | (+) |
| NHP2 | (-) | (+) | NT | (-) | NT |
| NHP3 | (-) | (-) | NT | (+) | NT |
| NHP4 | (+) | (+) | NT | (+) | NT |
| NHP5 | (+) | (-) | (+) | (+) | NT |
| NHP6 | (+) | (+) | NT | (+) | NT |
| NHP7 | (+) | (-) | NT | (-) | NT |
| NHP8 | (+) | (+) | NT | (+) | NT |
| NHP9 | (+) | (-) | NT | (+) | NT |
| Total of positives | 6 | 4 | 1 | 6 | 1 |

(+) = Positive (-) = Negative NT = not tested

**Table 2.**
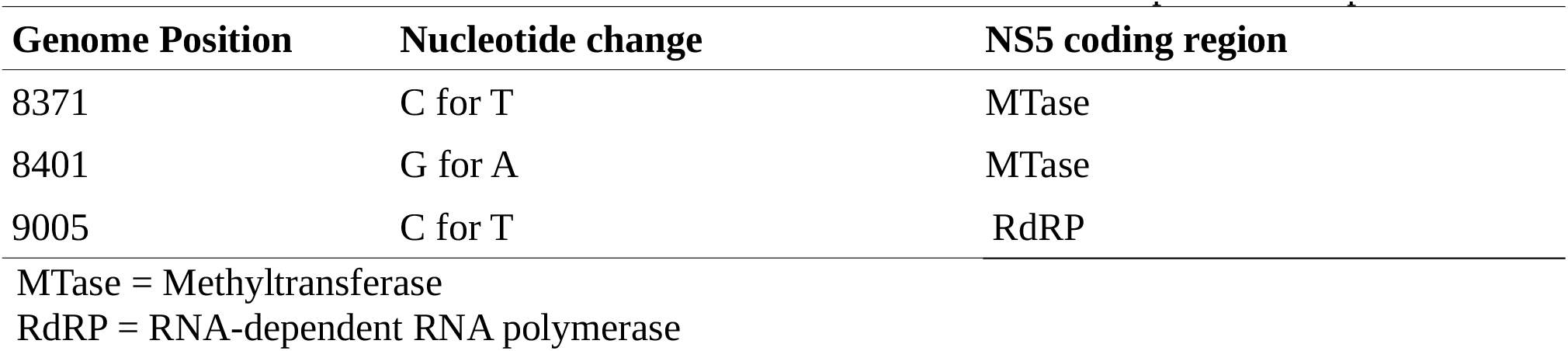
Nucleotide substitutions observed in *Flavivirus* nested-RT-PCR positive samples.

| Genome Position | Nucleotide change | NS5 coding region |
| --- | --- | --- |
| 8371 | C for T | MTase |
| 8401 | G for A | MTase |
| 9005 | C for T | RdRP |

MTase = Methyltransferase RdRP = RNA-dependent RNA polymerase

**Figure 1.**
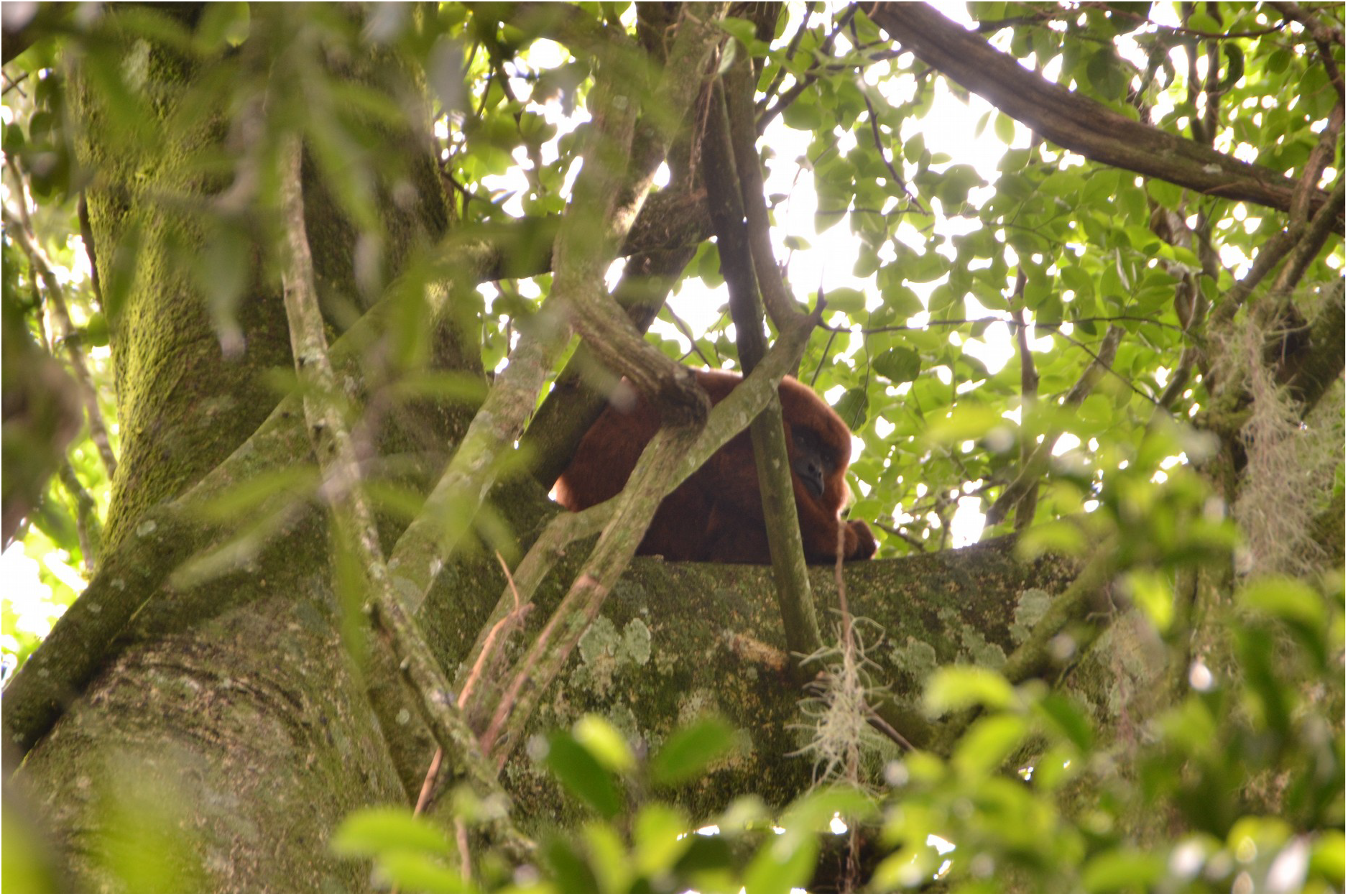
Free ranging healthy male howler monkey (*Alouatta guariba*) in an Atlantic Forest fragment.

**Figure 2.**
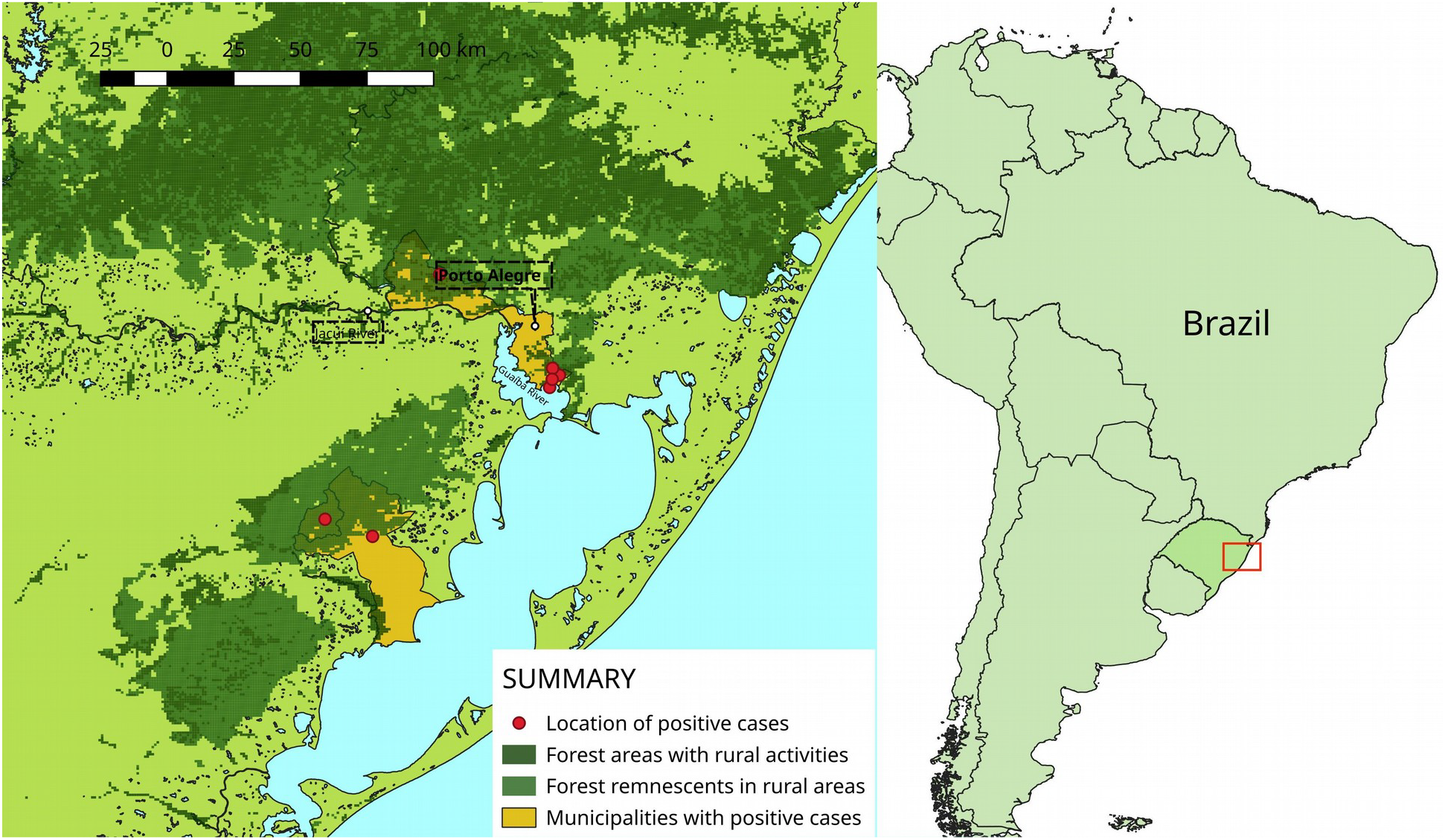
Rio Grande do Sul state, Brazil. Location of positive Flavivirus NHP, with four cases in Porto Alegre.

The resulting amplification products were purified and sequenced by the Sanger method. Amplification products ranged from 274 to 1007bp, and comparison with GenBank Databases revealed 96% to 98% identity with ZIKV lineage MR766.

### qRT-PCR Results

All nested RT-PCR positive samples underwent qRT-PCR and a neonate received after the outbreak described above (NHP10) that was negative in the nested RT-PCR screening but had lesions compatible with CZS was also included, and NHP 10 was positive in the liver, with a CT value of 34,61.

### Sequence analysis

Tree inference revealed clustering with African lineages of ZIKV (Figure 3). The best sequence obtained from the samples was submitted to GenBank database and received the accession number MK451708. Sequence alignment revealed consistent nucleotide substitutions in 3 positions of the alignment (genome positions 8372; 8402; 9005).

**Figure 3.**
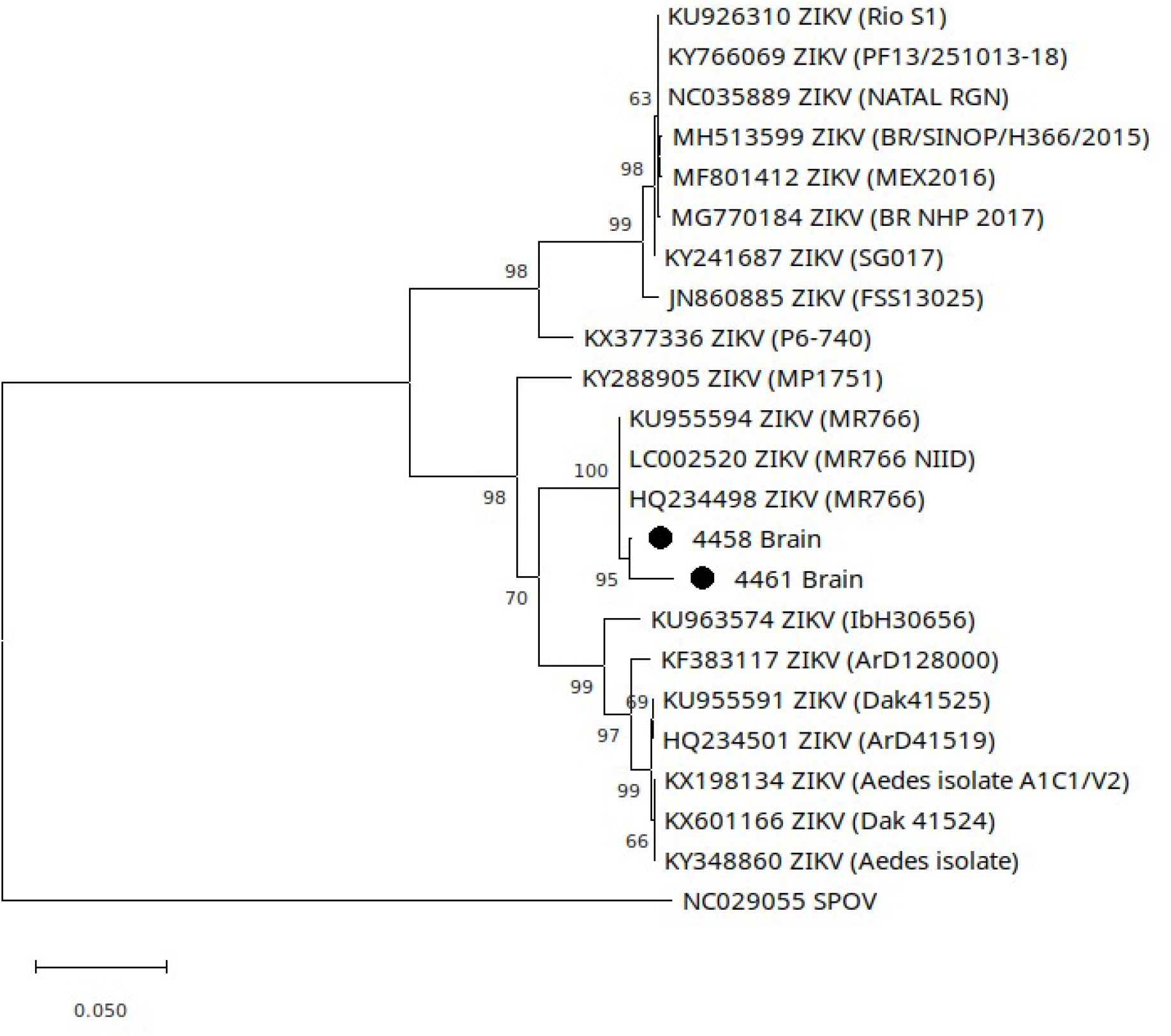
Molecular Phylogenetic analysis by Maximum Likelihood method using a dataset of ZIKV NS5 coding region, sequenced samples are labeled with a dot.

The tree using envelope sequencing data set (Figure 4) revealed close proximity to MR766 strain. Comparison of the 198bp overlapping region with the control revealed differences in 6 positions, with 4 substitutions and 1 insertion resulting in 1 glutamic acid addition in position 233 of the translated envelope sequence when compared to control and other ZIKV sequences aligned. Moreover, the envelope sequence presented absence of N-glicosilation in position 153-156 of the envelope translated sequence, similar to low passage strains of ZIKV (Figure 5).

**Figure 4.**
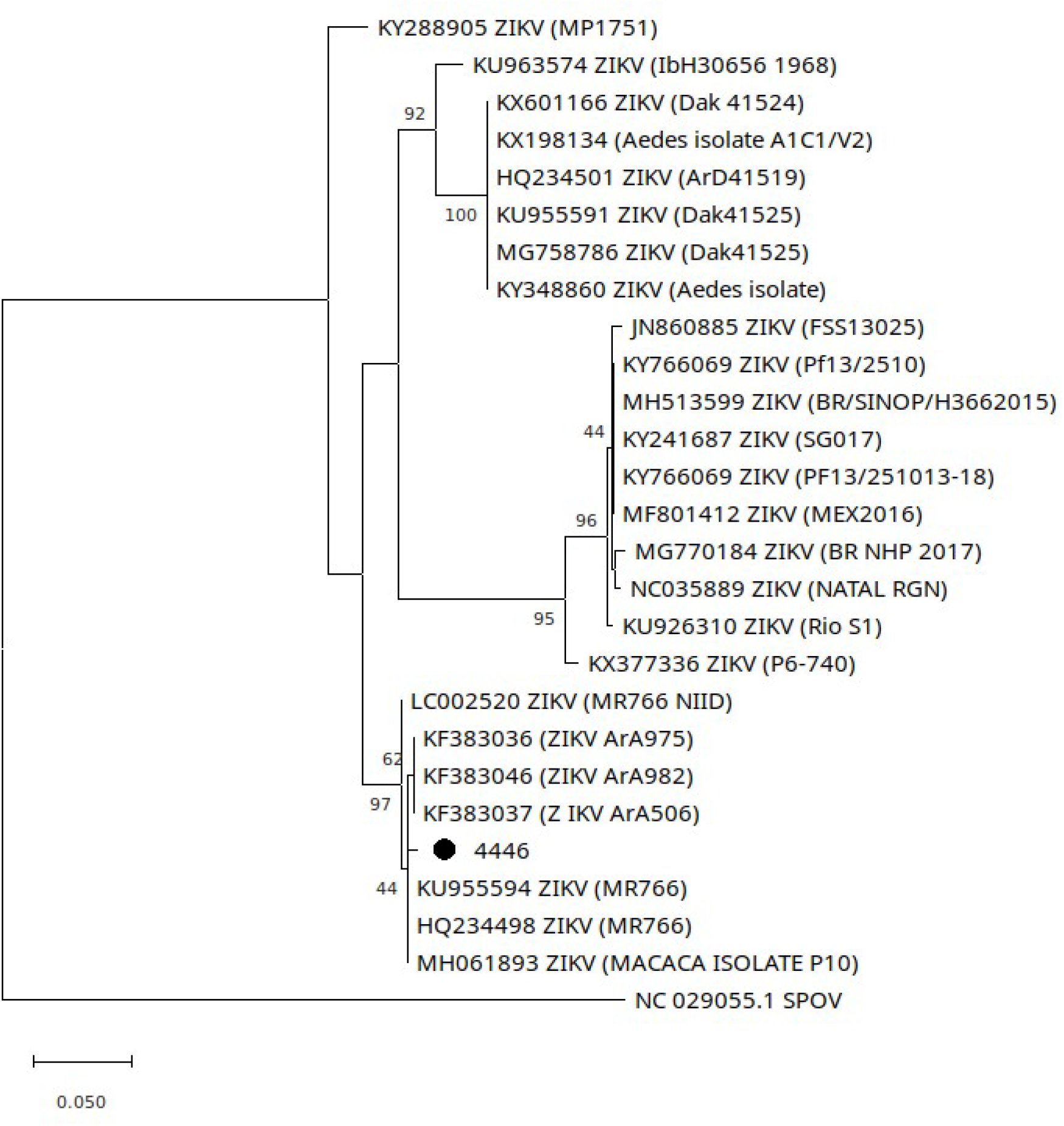
Molecular Phylogenetic analysis by Maximum Likelihood method using a dataset of ZIKV envelope coding region, sequenced sample is labeled with a dot.

**Figure 5.**
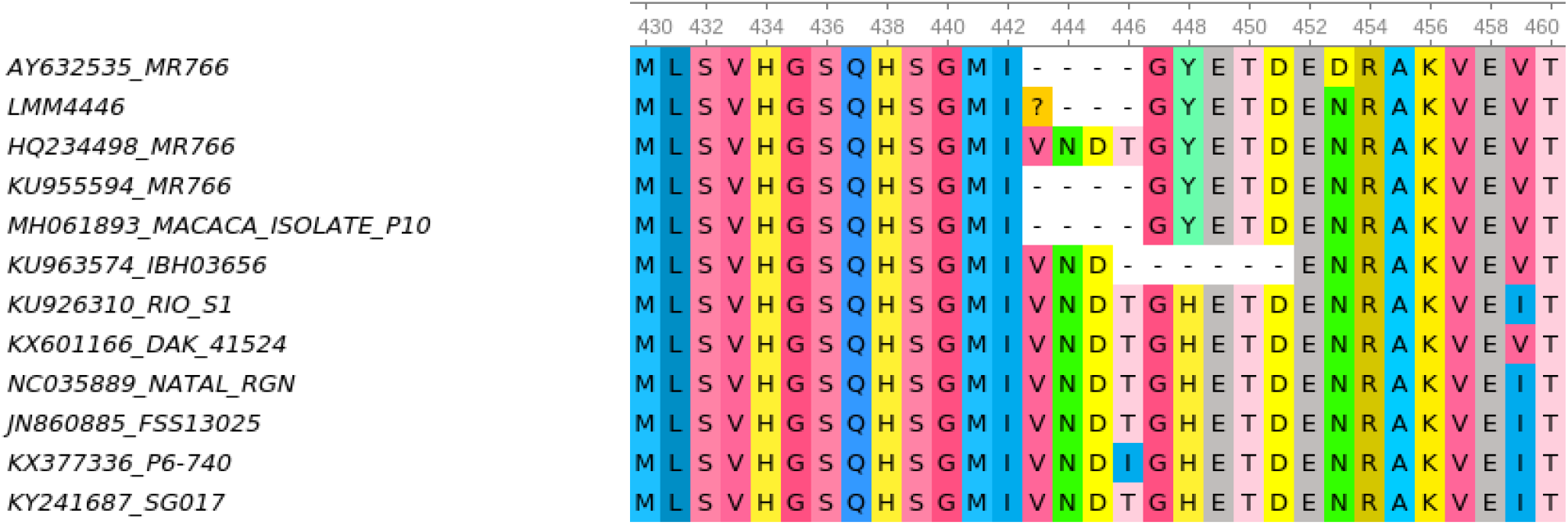
Translated envelope sequences showing N-glicosilation site at position 153-156. The sequenced sample (LMM4446) did not present the N-glicosilation site at position 153-156 of the envelope deduced sequence.

### Pathology and Immunohistochemistry

All positive NHP were *Allouata guariba* individuals, and 4 of them belonged to the same city and were submitted for necropsy during the same week. Eight of the positive NHP suffered trauma. Microscopically, *Flavivirus* positive NHP individuals presented inflammatory infiltrates in several degrees, specially in liver and kidney (Figure 6). Main lesions unrelated to other specific infections affecting positive NHP are listed in Table 3.

**Table 3.**
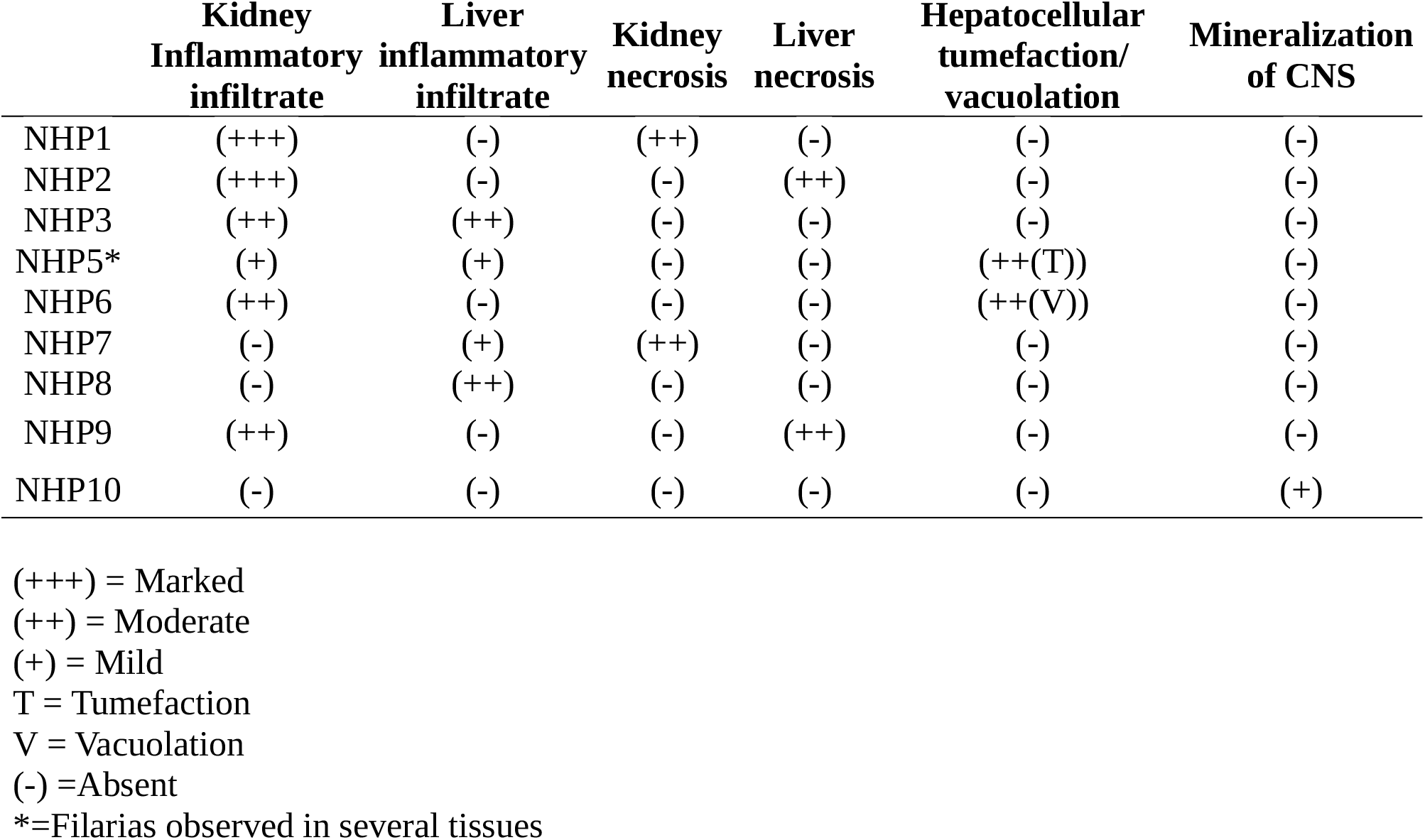
Main histologic features unrelated to other agents observed in *Flavivirus*-positive NHP.

**Figure 6.**
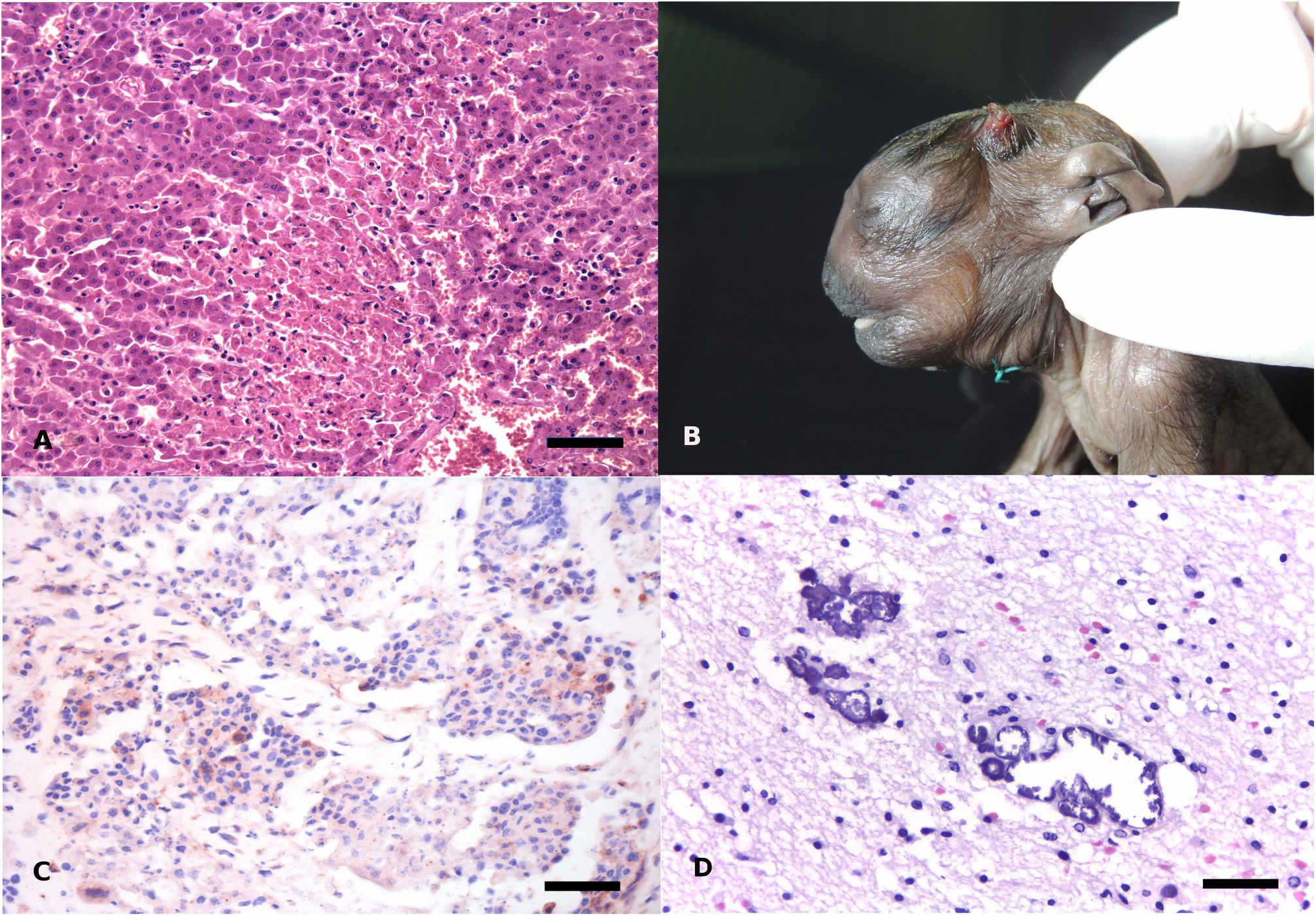
Tissues from several positive *Alouatta guariba* presented here, with histopathological changes and IHC labeling. A. Liver (NHP2) presenting marked coagulation necrosis, HE, Scale bar= 110µm. B. NHP10 with disproportional craniofacial sizes and deffective cranial closure. C. Placenta (NHP1) with ZIKV positive cells, IHC (Hematoxylin-AEC), Scale bar = 89µm. D. Brain of neonatal *A. guariba* (NHP10) presenting areas of mineralization, HE, Scale bar = 53µm.

NHP10 was received in Veterinary Pathology Sector of the Federal University of Rio Grande do Sul one year after NHP9. NHP10 belonged to a free ranging group. Grossly, the neonate presented opened sutures of cranial bones and absence of left frontal bone, excessive nucal skin, craniofacial disproportion (Figure 6, B) and palatoschisis. Lung flotation indicated that the animal breath, therefore was not a stillbirth.

Histologically, multiple mineralization foci were observed scattered throughout white matter in the brain, other tissues did not present significant alterations (Figure 6, D).

NHP4 could not be properly histologically evaluated due to autolysis; NHP5 presented *Filaria* sp. in several tissues, often with inflammatory infiltrate associated. Acute inflammation, hemorrhage, edema and aspiration pneumonia were observed in lungs from 3 cases of trauma, therefore are not listed in Table 3.

Lymphocytes infiltrating placental tissue presented ZIKV NS1 positive granular IHC staining in the cytoplasm (Figure 6), tissues from the fetus of this female were negative. NHP4 was positive in bile duct epithelial cells and renal tubular epithelial cells (Table 4 and Figure 6).

**Table 4.** IHC results per tissue in *Flavivirus* nested-RT-PCR positive NHP.

|  | CNS | Liver | Kidney | Placenta | Spleen |
| --- | --- | --- | --- | --- | --- |
| NHP1* | (-) | (-) | (-) | (+) | (-) |
| NHP2 | (-) | (-) | (-) | NT | (-) |
| NHP3 | (-) | (-) | (-) | NT | (-) |
| NHP4 | (-) | (-) | (-) | NT | (-) |
| NHP5 | (-) | (-) | (-) | NT | (-) |
| NHP6 | (-) | (-) | (-) | NT | (-) |
| NHP7 | (-) | (-) | (-) | NT | (-) |
| NHP8 | (-) | (-) | (-) | NT | (-) |
| NHP9 | (-) | (-) | (-) | NT | (-) |
| NHP10 | (-) | (-) | (-) | NT | (-) |

(+) = Positive (-) = Negative NT = Not tested \*Fetus from NHP1 was negative in the IHC test and was not tested with nested RT-PCR

## Discussion

The detection of this ZIKV-like *Flavivirus* in NHP living in the Atlantic forest remains from Southern Brazil emphasizes the adaption of this species to NHP. Since the discovery in 1947, ZIKV from lineage MR766 was only isolated from NHP and forest dwelling mosquitoes in Africa^6^. The sylvatic establishment of ZIKV in the Americas is among the major concerns for human health nowadays ^5^. The occurrence of an African lineage ZIKV in Southern Brazil, where the overall prevalence of Arbovirus infections in humans is believed to be low, elevates the importance of investigating circulating viruses with potential to adapt to human populations, since the cases described here are not a spillback event from the Asian-related ZIKV currently circulating in humans.

*Alouatta guariba* is highly susceptible to YFV, serving as a sentinel species indicating circulation of YFV ^17,18^. The susceptibility of this species to ZIKV has not been described to date. History of trauma was evidenced 8 out of 9 NHP, physical injuries to NHP are extremely difficult to occur, unless they are habituated to humans and other animals, or severely affected by an infection. All NHP presented here were free ranging animals, therefore the latter hypothesis is more likely, especially after the detection of this *Flavivirus*.

The most remarkable lesions observed were affecting kidney and liver. Inflamatory infiltrates and viral persistence in several tissues are described after ZIKV experimental infection ^14,15^. ZIKV induces apoptosis in kidney cells and lymphocytic infiltration in infected mice, and African-lineage ZIKV is better adapted to human kidney cells than Asian-lineage ZIKV^14^. NHP studied here did not present other causes for the infiltrates and necrosis observed in kidney tissue, nonetheless Mild inflammatory infiltrates without a defined cause can occur in kidney, specially in free ranging NHP, therefore it is difficult to draw conclusions regarding the histopathological findings in the kidneys. ZIKV leads to inflamatory infiltrate in several tissues, experimental infection in Rhesus Monkeys with Asian ZIKV induces inflammatory infiltrates and viral persistency in liver^16^. In this study, NHP presented inflammatory infiltrates in several degrees, degeneration and necrosis in the liver. NHP2 was PCR-positive in the liver and presented diffuse marked coagulation necrosis and hemorrhage. Hepatocyte necrosis and degeneration in NHP should be further investigated in order to understand if the virus detected in this study is involved in the pathogenesis of these histological features. CZS is characterized by developmental disturbances observed in several organs, including microcephaly, brain malformations, cranial deffects, ocular malformations, arthrogriposis and contractural malformations ^19^. Histologically, calcifications are the commonest feature observed in ZIKV infected neonates ^10^. NHP10 presented all lesions compatible with ZIKV infection, and was a free ranging animal, indicating that there is a degree of ZIKV circulation in sylvatic environments of Southern Brazil. This howler was found in the Lami reserve, so it is likely that the ZIKV detected belonged to the same lineage that was detected in all others, since it has been shown that African ZIKV can cause congenital CNS malformations in experimentally infected mice^20^, however, sequencing must be performed in order to conclude this statement.

Granular, perinuclear ZIKV positive IHC labeling was observed in lymphocyte infiltrates surrounding necrotic tissue in the placenta in NHP1(Figure 6). ZIKV persists in several tissues, including kidney, lymph nodes, placenta and spleen and also elicits inflamatory infiltrates in multiple organs ^13,15,16^, therefore lymphoid cells are likely to have viral antigens. ZIKV infection in placenta tends to be focal instead of diffuse and widespread^13^. The absence of IHC labeling in additional tissues can be due to immunological clearance and very focal infection^15^. The negative fetus in the case where placenta was positive (NHP1) can also be attributed to focal infections, assay sensitivity, or viral clearance^13^. Also, the capability of this strain to infect fetus passing placental barrier is unknown and might be lower than the capability of the Asian lineage to cause fetal infection. The labeling observed was consistent to previously described ZIKV IHC regarding location and shape^21^, the antibody utilized targets ZIKV NS1. NS1 detection in serum is a diagnostic marker of *Flavivirus* infection^22^, this protein is involved in immune evasion and pathogenicity, and the antibody used in this study presents no cross reactivity with other flaviviruses^21^.

NS5 fragment sequencing revealed 3 mutations of which, 2 were located in the MTase coding region and 1 was in the RdRP coding region. Unlike Asian lineages, ZIKV African lineage MR 766 has a highly conserved genome among isolates, with 0,4% nucleotide divergence in the entire genome, this feature could be responsible by the low pathogenicity of this lineage for humans^6^.

RdRP functioning activity motifs are highly conserved in ZIKV genome^22^. The sequence analyzed included three of these motifs, therefore, it was expected to observe few nucleotide substitutions. To date, African isolates were usually isolated from NHP and mosquitoes, whereas Asian ZIKV lineages were isolated from humans and mosquitoes, this demonstrates the high level of adaption of the African lineages to NHP host^6^. Isolated events of intrahost single nucleotide variations were observed in four samples, 3 of them were substitutions of A for G and one was a substitution of T for C. One of the samples presented an GAG insertion in 9037 to 9040 position of the genome. Intrahost single nucleotide substitutions are expected in RNA viruses^23^. The envelope sequencing comparison to the positive ZIKV MR766 control used in the lab revealed 6 differences in 198bp. This finding, along with the fact that extreme precaution was taken in order to prevent contamination, and IHC was performed 40km away from where PCR and viral isolation occurred, indicates that samples were not accidentally contaminated. All PCR reactions were repeated without positive control, from RNA extraction to electrophoresis, in order to confirm positive results.

The threat of a sylvatic cycle establishment of flaviviruses has been evidenced in many scientific studies^5, 7,8^. ZIKV has been demonstrated in NHP in southeastern Brazil, however, the source of this cycle involving marmosets was the human ZIKV epidemic event that started around 2013, as demonstrated by the phylogenetic characterization of this strain^8^. Conversely, the ZIKV strain detected here apparently is closer to African-lineage ZIKV than American lineage ZIKV, nonetheless, further sequencing is needed to fully understand the relation of this lineage with the currently known ZIKV lineages. Regardless of the phylogenetic classification of this ZIKV, results presented here highlight a formerly unknown disease affecting howler monkeys in Brazil.

Moreover, the effect of the circulation of this agent in people should be investigated, as there was a very low prevalence of ZIKV related autochtonous congenital syndrome cases in RS state^24^.

The origin of the virus reported here is uncertain and unless three main hypothesis are considered. The first is that this virus would have gone through cospeciation with its primate host. The origin of primates from the order Platyrrhini is believed to date from approximately 30 million years ago^25^.

These animals arrived in South America and expanded their distribution southwards to southern Patagonia during the Miocen Climate Optima (MCO) and retracted with the rising of the Andes and the global cooling that occurred subsequently^26^. Since RNA viruses have high rates of mutation, and the strains circulating nowadays are probably the same as 50000 to 100000 years ago^27^, it is also unlikely that this cycle would remain unnoticed in nature without prior detection.

The second scenario is that this virus has been imported by the same time YFV during slavetrade and remained infecting NHPs silently. African lineage of ZIKV is genetically more stable than the Asian lineage^6^. ZIKV circulating today has emerged in nature at approximately 100-130 years ago^3^, therefore it is possible that a common ancestor of the current African ZIKV has been brought to South America and due to its high adaptability to NHP and forest dwelling mosquitoes, established a sylvatic cycle that could remain silent due to an unrecognized syndrome affecting only NHP that tend to be vulnerable to flaviviral infections. The winter periods could have maintaned low replication rates and consequently low nucleotide substitution rates for this virus^27^.

A third hypothesis is that the virus was brought through human activities involving african animal importation, such as circus and zoo facilities. There are several zoo in RS, including in Porto Alegre metropolitan region with african NHP individuals, moreover, itinerating circus were common in Porto Alegre during the past centuries, as this city was in the circus route from Rio de Janeiro to Buenos Aires in the 19^th^ century ^28^.

The detection of this virus highlights the importance of improving surveillance strategies in sylvatic environments, since ZIKV is highly adapted to primates and mutations occurring fast in RNA viruses can promote a new disease emergency. Also, the detection of this virus can add knowledge regarding phylogenetic, prophylatic and therapeutic studies to collaborate with the understanding of the highly prevalent ZIKV circulating currently in people.

## Methods

### Sampling

In this study, tissues from 50 non-human primates (NHP) living in several regions of Rio Grande do Sul (RS) State, belonging to genus *Alouatta* sp., *Sapajus* sp. and *Callithrix* sp., received in the Veterinary Pathology Sector of the Federal University of Rio Grande do Sul from 2016 to 2018 were screened by nested Reverse Transcriptase Polymerase Chain Reaction (RT-PCR) for Flavivirus (Table 1). Pathological findings from positive NHPs were described, and virus isolation was attempted from these samples. *Flavivirus* positive tissue and isolation samples underwent Sanger sequencing in order to further characterize and classify flaviviruses found.

Samples were collected and fixed in 10% formalin solution for histological analysis. *Flavivirus* assay through RT-PCR, sequencing and viral isolation in cell culture were performed in frozen tissue samples from these NHPs.

### Nested RT-PCR

Frozen tissue specimens underwent nested RT-PCR applied for *Flavivirus* detection. RNA was extracted using TRIzol manufacturer instructions. Reverse transcription occurred using Promega GoScript™ cDNA synthesis kit, according to manufacturer instructions.

PCR reactions were performed using the Promega Colorless™ PCR kit. The first PCR reaction was performed as described by Bronzoni at al., 2005 ^29^, amplifying a 985bp fragment. For the second reaction, the primer set described by Moureau et al., 2007 ^30^ was applied in the PCR products of the first reaction. The second reaction resulted in an inner fragment of 274bp belonging to the non structural protein 5 (NS5) of the Flavivirus genome.

Amplification products underwent electrophoresis in a 2% agarose gel after nested RT-PCR reactions. The resulting products of the positive samples were sequenced and the sequence was searched in the BLAST platform in order to identify the sequences detected.

### qRT-PCR

All Flavivirus positive samples underwent qRT-PCR for ZIKV according to Lanciotti et al., 2008. Additionally a neonate sample that resulted negative in the nested RT-PCR, received after the outbreak, underwend qRT-PCR.

### Sequence Analysis and Phylogenetic Tree Inference

Positive sequences were aligned with NS5 of ZIKV MR766 sequences obtained in GenBank (Accession numbers: AY632535.2; DQ859059.1; KU720415.1; KU963573.2; KY989511.1; MH061859.1; MH061860.1; MH061875.1; MH061877.1; MH061878.1; MH061884.1; MH061885.1; MH061887.1; MH061888.1; MH061901.1; MH061904.1; MH061909.1; MH061910.1; MH0691911.1; MH061912.1; MH061913.1; MH061914.1; MH061915.1; MH130094.1; MH130095.1; MH130096.1; MH130107.1; MH130108.1; MH130109.1; NC012532.1) using Clustal Omega in the UGENE package for comparison of nucleotide changes. Translated positive envelope sequence was compared to a data set of ZIKV envelope sequences belonging to different lineages and with different passage histories (Accession numbers: AY632535; HQ234498; KU955594; KU963574; KU926310; MH061893; KX601166; NC035889; JN860885; KX377336; KY241687), results are shown in figures 5 and 6.

Next, two trees were inferred using MEGA X software, one for the NS5 coding region and a second tree for the envelope coding region. The NS5 tree was inferred using two positive samples with wider sequenced regions and the following 823bp-long ZIKV sequences (Accession numbers:KU926310; KU955591; KU955594; KU963574 ZIKV; KX198134; KX377336; KX601166; KY241687; KY348860; KY766069; KY288905; LC002520; MF801412; MG770184; MH513599; NC035889; KF383117; JN860885; HQ234501; HQ234498) and a Spondweni virus (accession number: NC029055.1) NS5 sequence as outgroup. A tree was inferred in MEGA X software using the Maximum Likelihood method based on the General Time Reversible model (GTR) ^31, 32^. The pylogenetic tree based on a fragment of 367bp of the envelope sequences using the sequence of one positive sample was inferred with the following ZIKV sequence data set: KU926310, KY241687, KY766069, MH513599, NC035889, MF801412, MG770184, JN860885, KF383035, KF383036, KF383037, KF383046, KX377336, KU955591, MG758786, HQ234501, KX198134, KX601166, KY348860, KY288905, KU963574, KU955594, LC002520, HQ234498, and the same outgroup as in the NS5 tree. MEGA X software was used for tree inference, using the Maximum Likelihood method based on GTR^31, 32^.

Additionally, samples underwent RT-PCR aiming two fragments of approximately 500bp, which overlapped in 198bp and the resulting products were sequenced by Sanger method. Positive control product from one of these RT-PCR underwent sequencing to discard possibility of contamination. It was possible to obtain a good quality contig from one sample, assembled in triplicates, from the second amplification region. The overlapping region of 198bp was compared to control sequence.

### Histopathology

After fixation in 10% formalin solution and paraffin embedding, the samples were cut in 3-5 micrometer thick sections and stained with hematoxylin and eosin^33^. Sections of brain, cerebelum, lung, liver, kidney, heart, stomach, jejunum and ileum, colon and cecum, adrenal gland, skeletal muscle, spleen, placenta and fetus (when present) were examined.

### Immunohistochemistry

Sections of brain, cerebellum, liver, placenta (when present) and kidney underwent IHC staining using MACH4 polymer. Endogenous peroxidase blocking was performed through incubation of the sections in a 3% hydrogen peroxide in methanol solution for 20 min. Citrate buffer (pH 6.0) was applied for antigen recovery at 96ºC for 40 min. Slides were incubated in milk diluted in phosphate-buffered saline (PBS) for 20 min in order to block unspecific reactions.

Monoclonal antibody against ZIKV Non-Structural 1 (NS1) protein (Arigo Biolaboratories, Taiwan, Republic of China) was diluted 1:1000 in (PBS) and applied to the slides in a humid chamber overnight at 4ºC. Subsequently, slides were incubated with Mach 4™ Universal AP Polymer Kit system (Biocare Medical) according to manufacturer recommendations and stained with AEC chromogen (Biocare Medical, USA). Next, slides were counterstained using Mayer hematoxylin and mounted routinely. Vero cell culture inoculated with ZIKV MR766 was used as positive control, the same cell culture without inoculus was utilized for negative control. Additionally, histologic sections of the positive control incubated with PBS were used as negative controls.

